# Behavioral responses of household ants (Hymenoptera) to odor of different coffee species and formulations: Sustainability approach for green pest management strategies

**DOI:** 10.1101/101303

**Authors:** Abdul Hafiz Ab Majid, Hamady Dieng, Siti Salbiah Ellias, Tomomitsu Satho

**Affiliations:** Household and Structural Urban Entomology Laboratory, Vector Control Research Unit, School of Biological Sciences, Universiti Sains Malaysia; Institute of Biodiversity and Environmental Conservation, Universiti Malaysia Sarawak, 94300 Kota Samarahan, Sarawak, Malaysia; Department of Microbiology, Faculty of Pharmaceutical Sciences, Fukuoka University, 8-19-1 Nanakuma, Johan-ku, 814- 0180 Fukuoka, Japan

**Keywords:** household ants, olfactory, coffee, behavioral effects

## Abstract

Odor sensation is a sensory modality of considerable significance in the foraging behavior and interactional organization of ants. In the food bait technology, smell is the basis of attraction, which in turn, is the initial line of bait use and a key parameter for judging efficacy. Yet, baits that are currently available possess low attractiveness to many ant pests. Hence, strategies to produce ant bait with increased attractiveness are needed. Despite evidence that coffee has a diverse aroma complex that affects the behavior of honey bees, which share many social and genetic traits with ants, its attraction to formicine foragers have yet to be investigated. In a series of Y-tube olfactometer bioassays, we examined the behavioral responses of *Tapinoma indicum* (TI), *Monomorium pharaonis* (MP) and *Solenopsis geminata* (SG) to various coffee-induced odor stimuli, comprised of extracts from Arabica, Robusta and Liberica. All coffee species showed an influence on the behavior of TI, MP and SG workers, but the clearest effect was seen with Arabica. The workers of TI, MP and SG were more attracted to the odor of 0.01% Arabica extract (ONE), in comparison with 0.05% Arabica extract (TWO) or 0.05% Arabica extract (THREE). Arabica extract mixed with sugar (S) elicited significant attraction from workers of all three species when in balanced competition with either unsweetened Arabica extract or water. These results indicate that coffee, particularly Arabica is attractive to the foragers of TI, MP and SG, and can encourage the use of coffee in bait development. As coffee is already accepted in most human societies, the development of coffee-based bait may be an attractive option.

## INTRODUCTION

There are more than 12,000 known species of ants (Hammond, 2011), many of which are among the most common insects invading or living inside human establishments where they become a nuisance and cause damage (Lee, 2002; Coggins, 2013). This is typical of formicine ants such as *Tapinoma indicum*, regarded as household pests that invade health care facilities in Southeast Asia (Lee, 2002; Man and Lee, 2014); *Monomorium pharaonis,* a serious threats to homes, hospitals, offices and business establishments—hotels, supermarkets, restaurants (Osae, Cobblah, Djankpa, Lodoh, and Botwe, 2011); and *Solenopsis geminata*, a threat to birds, livestock, agricultural ecosystems and people (Harris, et al. 2005).

*T. indicum* has an increased ability to invade disturbed habitats as they are able to rapidly find suitable habitats for nesting and produce large-sized colonies (Passera, 1994). When they invade houses, *M. pharaonis* also known as the Pharoah ants, attack a wide range of foodstuff, clothes, books (Dumpert, 1981) and can chew on silk, rayon, rubber and electrical wiring (Hölldobler and Wilson, 1990). In hospitals, the Pharoah ants are reputed to carry several pathogenic bacteria (Haack and Granovsky, 1990; Smith and Whitman, 1992) and may feed on wounds (Anon, 1986), thus considered a potential disease vector (Osae, et ak. 2011). *S. geminata*, on the other hand, is a major threat to agricultural crops (Wilson, 2005), affecting farmers’ performance by inflicting irritating stings (Hill, 1987; Nestel and Dickschen, 1990). This type of ants also causes substantial damage to PVC coatings of electrical wiring (Prins, 1985) and drip irrigation tubing (Chang and Ota, 1990).

Efforts to combat such damages rely heavily on the use of chemical insecticides through baiting (Higgins, Bell, Silcox, and Hollbrook, 1997), residual perimeter sprays (Potter and Hillery, 2002), or both in combination (Higgins, Bell, Silcox, and Hollbrook, 1997). In these global strategies against ant pests, baiting form a very crucial part of the solution (Benett et al., 2013). This method takes advantage of the social trophallactic and grooming behaviors of ants (Lee, 2008) and relies on the pick up of bait particles by foraging workers and their transfers to other colony members (Benett et al., 2013). Baits have been used successfully to control a number of social insect pests (Stanley, 2004; Benett et al., 2013). However, many bait-based programs failed due to insecticide resistance and insufficient level of attractiveness (Rust, Reierson, and Klotz, 2002; Krushelnycky and Gillespie, 2008). Thus, the ability to produce bait with high attractiveness remains a major challenge for the bait-based ant control strategy. For target ants to pick up and transfer bait particles to the colony, they must be attracted to the bait. Therefore, the first and critical step in the bait use process is attracting the foragers (Benett et al., 2013).

The sense of smell is the main sensory modality for ants (Gronenberg, 2008). In fact, they primarily perceive the world through smell via the detection of airborne chemicals and touch (Dodgson, 2015). Since the first line in the bait use process is to entice foragers (Benett et al., 2013) and that olfactory cues are essential for social organization and behaviors (Gronenberg, 2008), especially orientation (Wolf and Wehner, 2000), these aspects of ant biology should receive the highest attention when seeking new control strategies. Despite substantial efforts to increase bait attractiveness by the addition of insect tissue (Williams, Lofgren, and Vander Meer, 1990), different fruit juices (Lucas and Invest, 1993), sugars, proteins, oils, or a combination of different attractants (Rust and Choe, 2012), very few baits are available that are very attractive to ants (Rust, Reierson, and Klotz, 2002; Krushelnycky and Gillespie, 2008). Therefore, it is an important strategy to target strong-smelling and chemically rich materials.

A special characteristic of ants is that they have a remarkably sophisticated sense of smell (Choe, Ramírez, and Tsutsui, 2012; Harris-Lovett, 2015) and seem to be able to sniff most of the substances that humans can smell (Dodgson, 2015). They have a brain with around 250,000 nerve cells (John and Sarah, 2010) and use special neurons associated with tiny hairs on their antennae to smell odors (Roces, 1994; Dupuy, et al., 2006). An ant’s antennal lobe possesses over 400 glomeruli (Zube, et al., 2008; Gronenberg, 2008). There is a close association between a number of glomeruli and the smell detection capacity. Indeed, a large number of odorant receptors and glomeruli allows for complex combinatorial olfactory codes to detect and discriminate a wide range of odorant stimuli (Kelber, et al. 2009) with precision (Gronenberg, 2008). Therefore, any material that has high aroma potency may be a valuable candidate as an attractant for ants.

There are more than 1000 chemicals in coffee including over 800 aromatic compounds (Clarke and Macrae, 2012). Many of these constituents such as aliphatics—carbonyl and sulfur contain compounds: alicyclic elements—ketones; aromatic benzenoid compounds—phenols; heterocyclic compounds—furans, hydrofurans, pyrroles, pyridines, quinolines, pyrazines, quinoxalines, indoles, thiophens, thiophenones, thiazoles and oxazoles are produced during the roasting process (Illy and Illy, 2015; Fisk, Kettle, Hofmeister, Virdie, and Kenny, 2012; Clarke, and Macrae, 2012). The increased diversity of compounds results in an aroma complex comprising fruity, earthy, catty, roasty, spicy, buttery, sweet, rotten cabbage-like, honey-like, potato-like, caramel-like, seasoning-like, and vanilla-like smells (Grosch, 1998). These smells hit many scent receptors in both humans and animals including insects. For instance, exposure of rats to coffee aroma deprived them of sleep for a day (Seo et al., 2008). In honey bees, caffeine from coffee stimulated responses in olfactory learning and memory (Wright et al., 2013). Many processes inherent to social living and food processing in bees possess homologs in ants. As bees are genetically related to ants (Johnson et al., 2013) and have far more glomeruli than them (160 glomeruli) (Flanagan and Mercer, 1989), they may be highly responsive to the aroma of coffee. Despite evidence that coffee has a variety of aromas and that coffee influences some hymenopterans (bees), there have yet to be any studies on the effects of coffee on ant foraging behaviors with respect to bait technology improvement. The present study was carried out to examine the behavioral responses of formicine ants towards extracts from different coffee species. The behaviors of these ants in response to coffee exposure at three different concentrations and sugar contents were also probed.

## MATERIALS AND METHODS

### Test ant species and experimental subjects

*T. indicum* (TI), *M. pharaonis* (MP) and *S*. *geminata* (SG) were selected in this study. For each of these ant species, colonies were located within the Minden Campus of University of Science Malaysia (Penang, Malaysia, latitudes 5°8'N - 5°3'N; longitudes 100°8'E - 100°32'E; Ahmad et al., 2006). For each of the three formicine ants, two different colonies were selected as sources of test subjects, and each was marked using tagged wooden stakes. Active workers were collected by placing twenty to thirty traps within each colony starting from 7 am. Each trap consisted of a 1.5-mL Eppendorf tube with the lower bottom removed and was interiorly lined with a thin layer of Fluon (Polytetrafluroethylene suspension), adopting Shirwaikar, Issac, and Malini, (2004) in their efforts to prevent the escape of trapped workers. A minute amount of peanut butter or honey held on a small piece of paper was placed inside all traps to serve as feed. After 3 hours of collection, the trapped workers were kept in 250-mL plastic containers lined with Fluon coat and labeled according to species, colony number, and date of sampling. The workers’ samples were brought to the insectarium of the School of Biological Sciences (Universiti Sains Malaysia. Subsamples were kept in 90% ethanol for species identity confirmation. The laboratory conditions were 27°C ± 2.0°C, 75% ± 1% relative humidity (RH), in photoperiod 13:10 hours (light:dark) with 1 hour of dusk. For all species, workers that had acclimatized to the laboratory environment and were active after a 12h-starvation period, were used as experimental subjects.

### Coffee materials and experimental extracts

*Coffea arabica* (Arabica)*, C. canephora* (Robusta) and *C. liberica* (Liberica) varieties cultivated in Malaysia were selected for this study. Samples of 150 g of dried roasted coffee bean seeds from these different species were individually crushed using a blender (Pensonic Blender PEN-PB3103; Senheng^®^ Electric Sdn. Bhd., Kuala Lumpur, Malaysia) for 10 minutes. In order to obtain uniform textures, grounds were sieved twice through a kitchen sieve strainer (250-wire mesh). Fine grounds were used to produce different extracts by Soxhlet extraction in accordance with the published procedures (Cholakov, Toteva, Nikolov, Uzunova, and Yanev, 2013). Briefly, an amount of 50 g of fine Arabica grounds was placed on a paper thimble and extracted with 250 mL of methanol. Five hours after the first down-pouring of the methanol from the thimble, the resulting mixture (methanol + extract) was siphoned into a flat-bottomed flask, filtered and transferred into a glass petri dish. The extract was placed in an incubator (Memmert GmbH + Co, KG, Germany) set at 50°C. After 3 days of evaporation, the extract which was kept in a vial, stored in a fridge (-4°C) can validly be referred to as Arabica extract. The same quantity of fine grounds and procedures delineated above for Arabica were also performed for Robusta and Liberica grounds; the two resulting juices were designated as Robusta and Liberica extracts correspondingly.

Three other experimental concentrations of coffee were produced using Arabica, following a slightly modified method previously published (Derraik and Stanley 2005). An amount of 0.005 g of powdered roasted grounds was placed into a 150-mL glass cup with 50 mL of boiling water. After ten minutes, the resulting solution was sieved through a piece of fine-mesh mosquito net as done in other experiments (Satho et al., 2015); this 0.01% Arabica solution was designated as “ONE”. Solutions generated by submerging amounts of 0.025 and 0.05 g of Arabica grounds each in 50 mL of boiling water for the same extraction time (10 minutes) were referred to as Arabica extracts TWO (0.05%) and THREE (0.10%) respectively. To acquire a sugared experimental Arabica, we proceeded as follows. An amount 0.005 g of powdered roasted Arabica grounds was immersed in 50 mL of boiling water in a 150-mL glass cup and allowed to disintegrate. After ten minutes, the resulting solution was filtered, and 0.25 g of sugar was added to it. The infusion collected after complete dissolution of the sugar and filtration was referred to as Arabica extract with sugar or “S” (0.50% of sugar). Another solution obtained by the same procedures described for the above infusions, but without sugar is dubbed Arabica extract without sugar or “NS”. Cool boiled water acted as a control (C). When necessary, additional extract preparations were carried out to obtain enough quantities of experimental extracts for the bioassays.

### Bioassays setup

A Y-tube olfactometer system that has been described elsewhere (Yusuf, Crewe, and Pirk, 2014) was used to test the attraction of workers of different formicine ant pests to odors emanating from different coffee extracts. The system comprised a central tube (7.5 cm long) that branches into two lateral arms (each 5.75 cm long and 10 mm wide). Each arm was connected to a glass chamber (200-mL capacity) that held a given test coffee extract. Each glass chamber had a screw top containing inlets for the incoming air and outlets for odors to exit the Y-tube. A charcoal-filtered air stream was passed into each arm at a flow rate of 250 mL/min to drive experimental odors towards the decision area. A mesh screen was placed at each of the end points of the olfactometer to prevent the escape of workers from the test arena. Bioassays were conducted under laboratory conditions (temperature: 27.0 ± 0.1◦C, relative humidity: 80.5 ± 8.5%, and photoperiod: 14:10 h) between 7 am and 10 am or 4 and 5 pm, adopting the practice of previous researchers (Yusuf, et al., 2014) in their attempts to cover the optimal foraging periods of test ants.

A first experiment was carried out to examine the attractiveness of extracts from three coffee species to *T. indicum*. To determine whether TI is attracted to Arabica in the presence of Robusta, thirty workers were introduced individually into the central arm and given 5 min to choose between the stimuli in the two arms, each holding either Arabica or Robusta extracts. The same procedures and numbers of TI worker replicates as reported above were repeated but the two arms contained one the following solutions: (i) Arabica and (ii) Liberica. To establish whether TI was enticed by Robusta in the presence of Liberica, thirty TI worker replicates were individually given the opportunity to choose between two stimuli: (i) an arm holding Robusta and (ii) an arm containing Liberica. The same settings, treatments, procedures, and numbers of replicates, were also carried out for MP, and SG.

### Bioassay 2

To determine whether coffee attractiveness to ants depends on coffee concentration, the behavioral responses of TI workers to Arabica extracts were evaluated for three different concentrations in a series of two-choice comparative trials. In the first trial, a single TI worker was released from the central arm of the Y-tube and was allowed to select one of the following conditions: (i) an arm holding 0.01% Arabica extract (ONE) and (ii) an arm holding 0.05% Arabica extract (TWO). In the second trial, an individual TI worker was given a choice between (i) an arm holding ONE and (ii) an arm holding 0.10% Arabica extract (THREE). In the third trial, a single TI worker was given opportunity to choose between an arm holding TWO and the other with THREE. For each trial, thirty TI workers were individually tested and in each case, a single worker was given 5 min to select between the two treated arms. The same three comparative trials performed for TI were also done for MP and SG, with thirty test worker replicates in each case.

### Bioassay 3

To establish whether sugar has an influence on the attractiveness potential of coffee on ants, the behavioral responses of TI workers to three Arabica extracts with different sugar contents were tested in a sequence of two-choice comparative bioassays, using similar designs and procedures mentioned in Experiment 2. In a first bioassay, an individual TI worker was introduced into the central arm and permitted to choose between the two arms each containing one of the following Arabica extracts: (i) Arabica extract mixed with 0.50% of sugar (S) and ii) Arabica extract without sugar (NS). In the second bioassay, a single TI worker was given a choice between (i) an arm containing S and (ii) an arm holding water (Control: C). In the third bioassay, a single TI worker had to choose between an arm containing NS and an arm with C. Thirty TI workers were tested for each of the three bioassays. The same numbers of bioassays, worker replicates, procedures and treatments as described for TI were also conducted for MP and SG.

### Data collection and analyses

In all bioassays, each worker was given a 5-min period to select between the arms. A choice for the left or right arm of the olfactometer was recorded when the worker went 1 cm past the Y junction and stayed there for at least 1 minute or when it frequently visited an arm. For all two-choice bioassay data obtained, a preference index (PI) was computed using a formula reported by other researchers (Carsson et al., 1999; Nyasembe et al., 2012). The formula was expressed as PI = [(SS – NSS) / (SS + NSS)] x 100, where SS is the number of ants responding to the first stimulus and NSS is the number of them reacting to the second one. A PI of 0% would mean that equal number of experimental workers were attracted to each of the two arms of the Y-tube olfactometer. A positive value of the PI signifies that more workers responded to the first extract whereas a negative value indicates more were attracted to the second extract. As established by Anaclerio et al. (2012), in all bioassays, only workers that responded were included in data collection and analysis. The PI was used as a measure of the ants’ behavioral response to the tested stimulus. Behavioral responses of workers belonging to the different tested ant species to each experimental coffee stimulus were compared using the Chi-square test (SPSS Statistics, *v.* 20) with *P* < 0.05 as an indicator of statistical significance.

## RESULTS

### Behavioral responses to stimuli from different coffee species

When presented with Arabica and Robusta, TI is attracted to both odor stimuli, but Arabica seems to elicit more attraction from its workers. However, there is no significant difference in the preference index between the two smells (*χ*2 = 1.096, *df* = 1, *P* > 0.05). In the second experiment, a comparison is made between the attractiveness of the extracts of Arabica and Liberica. The TI workers tend to be more attracted to the odor stimulus from Arabica than that from Liberica, but the difference is not significant (*χ*2 = 2.25, *df* = 1, *P* > 0.05). When exposed to the smells from Robusta and Liberica, TI workers have a tendency to exhibit a preference for the first odor over the second one, but the attractiveness did not differ significantly between the coffee species (*χ*2 = 0.20, *df* = 1, *P*> 0.05) (Fig. 1A).

**Fig. 1.**
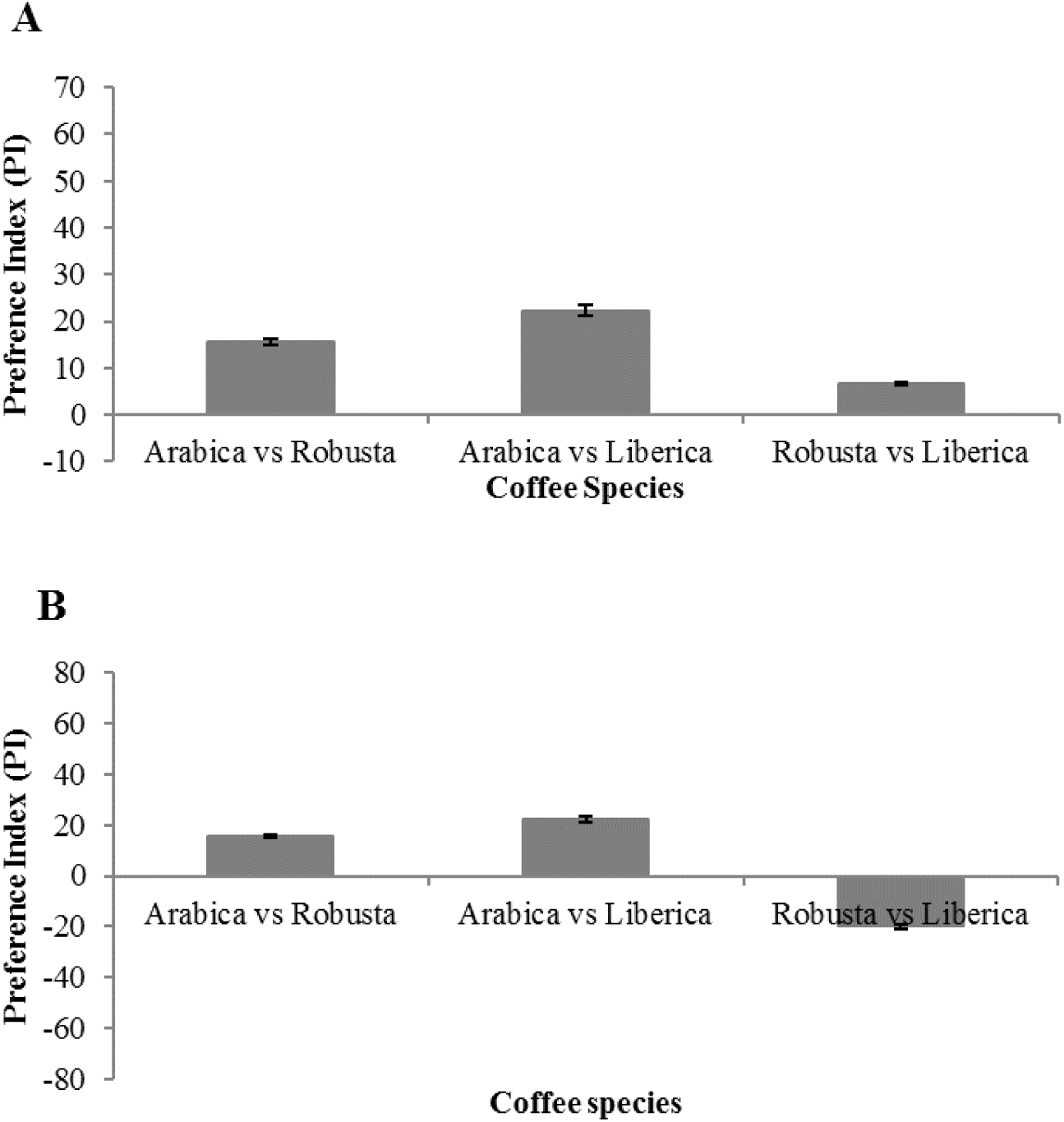
Responses of *T. indicum* (TI) (**A**), *M. pharaonis* (MP) (**B**), and *S. geminata* (SG) (**C**) in a Y-tube olfactometer where workers were given a choice between different binary combinations of odor stimuli from different coffee species. Positive bars indicate preference for the first extract. Negative bars indicate preference for the second extract. The asterisk over bars (∗) stands for significant difference (*P* < 0.05) within a dual choice test based on Chi square test.

When given similar options to walk towards the two sources of coffee odor, MP workers are attracted to both odors, but behavioral responses reveals different patterns according to the coffee type. Arabica has a tendency to attract more workers than Robusta, as shown by the positive PI, but the difference in attractiveness between the two coffee species is insignificant (*χ*2 = 1.096, *df* = 1, *P* > 0.05). When one arm holding Arabica and another with Liberica are presented to MP, they head to both odor sources, but the workers are more susceptible to approach Arabica than Liberica. However, the attractiveness level of the two coffee odor is statistically similar (*χ*2 = 2.25, *df* = 1, *P* > 0.05). In the choice bioassay involving Robusta and Liberica, the MP workers visit both coffee odor sources, but they tend to visit the arm holding Liberica more. However, there is no significant difference in the attractiveness level between the Robusta and Liberica stimuli (*χ*2 = 1.818, *df* = 1, *P* > 0.05), as illustrated by the negative PI (Fig. 1B).

When there is an equal chance for walking towards a source of coffee either the Arabica odor and Robusta smell, SG workers tend to be more attracted to the first odor. However, there are no significant differences in the attraction level between the two sources (*χ2* = 6.66, *df* = 1, *P* < 0.05). In the two-choice bioassay comprising Arabica and Liberica, SG workers walk towards both coffee odor sources, but they tend to prefer Arabica, but the attractiveness does not differ between the two odors (*χ2* = 2.25, *df* = 1, *P* > 0.05). There was no difference in the attractiveness level between Robusta and Liberica coffee (*χ2* = 0.20, *df* = 1, *P* > 0.05) (Fig. 1C).

### Behavioral responses following Arabica exposure at three different concentrations

In the two-choice experiment involving the two lowest Arabica concentrated extracts (ONE and TWO), TI workers are significantly attracted to extract ONE (*χ2* = 4.464, *df* = 1, *P* < 0.05). When presented with extract ONE and THREE, TI workers prefer the first over the second extract (*χ2* = 11.439, *df* = 1, *P* < 0.05). However, no significance in the level of attraction to TI workers is observed between TWO and THREE (*χ2* = 1.434, *df* = 1, *P* > 0.05) (Fig. 2A).

**Fig. 2.**
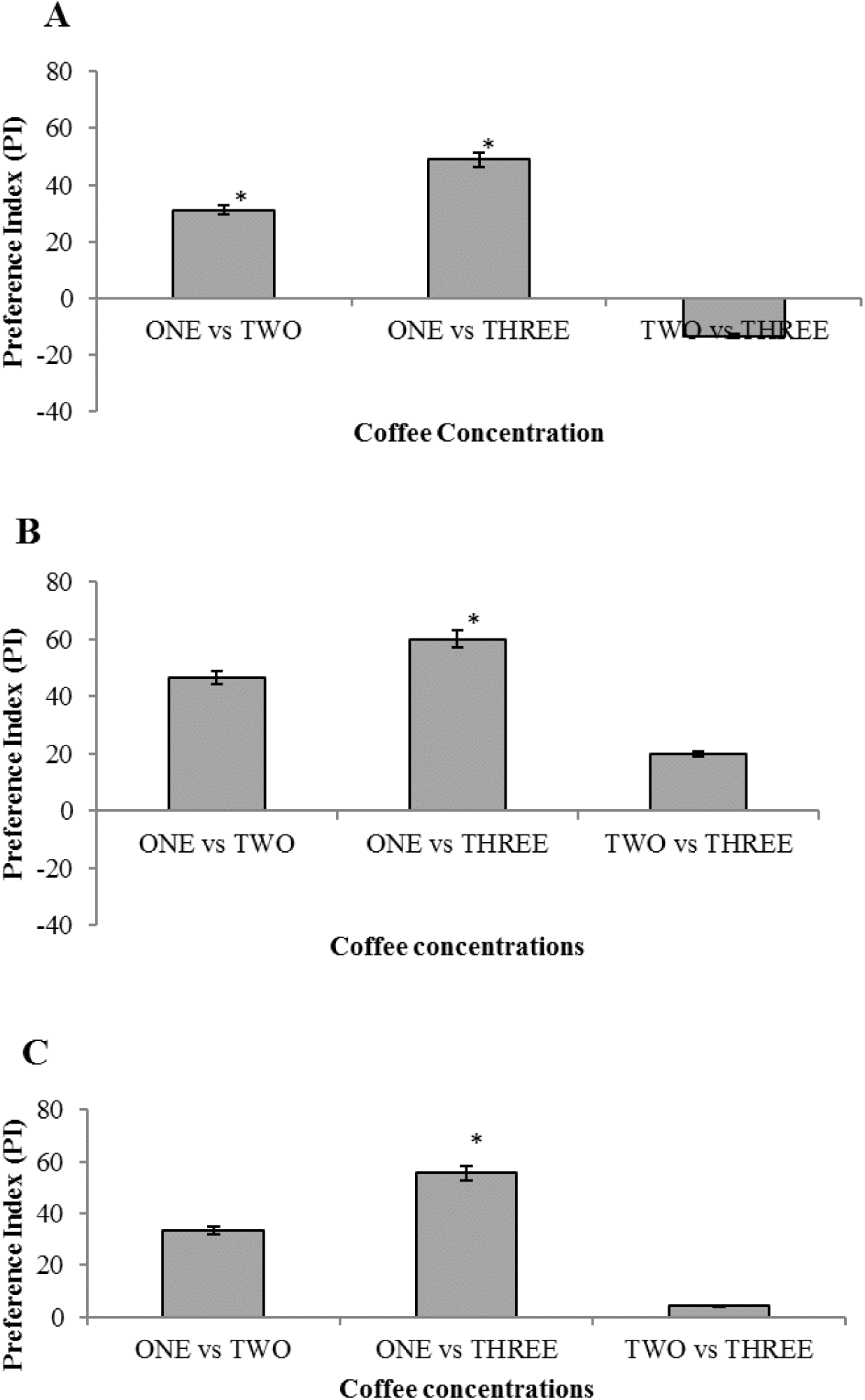
Responses of *T. indicum* (TI) (**A**), *M. pharaonis* (MP) (**B**), and *S. geminata* (SG) (**C**) in a Y-tube olfactometer where workers were given choices involving different binary combinations differently concentrated Arabica extracts. Positive bars indicate preference for the first extract. Negative bars indicate preference for the second extract. The asterisk over bars (∗) stands for significant difference (*P* < 0.05) within a two-choice test based on Chi square test. Arabica extract ONE (0.01%), Arabica extracts TWO (0.05%) and THREE (0.10%) respectively.

**Fig. 3.**
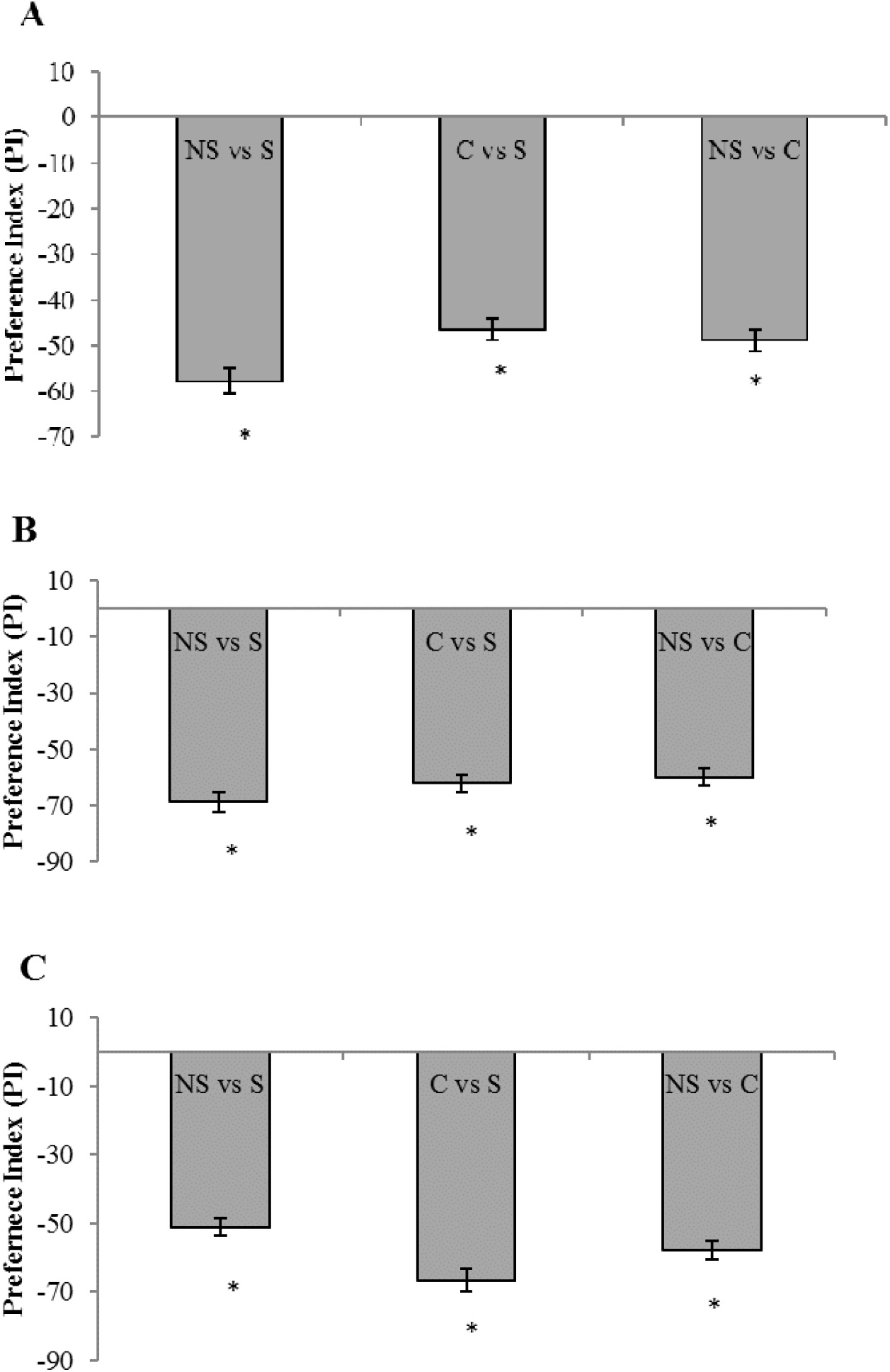
Responses of *T. indicum* (TI) (**A**), *M. pharaonis* (MP) (**B**), and *S. geminata* (SG) (**C**) in a Y-tube olfactometer where workers were given choices involving different binary combinations Arabica extracts with different sugar contents. Positive bars indicate preference for the first extract. Negative bars indicate preference for the second extract. The asterisk over bars (∗) stands for significant difference (*P* < 0.05) within a two-choice test based on Chi square test. S referred Arabica extract with sugar (0.50% of sugar); NS referred to Arabica extract without sugar; C referred to Cool boiled water acted as a control

When provided with similar chances to walk towards the Arabica ONE or the Arabica TWO, MP is attracted to both, but significantly more attracted to the less concentrated extract (*χ2* = 3.258, *df* = 1, *P* < 0.05). Similarly, Arabica ONE elicit greater attractiveness to MP workers when presented with Arabica THREE in a two-choice bioassay (*χ2* = 17.802, *df* = 1, *P*< 0.05). However, no significant attraction of MP is found when workers are furnished with even options to choose between two arms containing Arabica TWO and Arabica THREE (*χ2* = 1.818, *df* = 1, *P* > 0.05) (Fig. 2B).

No significant differences in the attraction to SG workers are found between Arabica ONE and TWO (*χ2* = 0.804, *df* = 1, *P* > 0.05), Arabica TWO and THREE (*χ2* = 15.05, *df* = 1, *P* < 0.05) or Arabica ONE and THREE (*χ2* = 0.089, *df* = 1, *P* > 0.05) (Fig. 2C).

### Behavioral responses following Arabica exposure at sugar contents

In the two-choice bioassay comprising the two less sweet Arabica extracts, Arabica S shows greater attractiveness to TI workers (*χ2* = 16.39, *df* = 1, *P* < 0.05). A similar pattern of preference is observed when comparing Arabica S and water (*χ2* = 20.854, *df* = 1, *P* < 0.05). A significant attraction of TI workers by water is noticed when they are given a choice between an arm containing Arabica NS and water (*χ2* = 11.439, *df* = 1, *P* < 0.05) (Fig. 3A).

Arabica S extract generated significant attraction in MP workers when in competition with Arabica NS (*χ2* = 24.23, *df* = 1, *P* < 0.05) and with water (*χ2* = 19.289, *df* = 1, *P* < 0.05). In the two-choice bioassay involving water and the sweetest Arabica extract, MP workers exhibited greater preference for water (*χ2* = 17.802, *df* = 1, *P* < 0.05) (Fig. 3B).

In the two-choice experiment, both Arabica extracts (S and NS) were attractive to SG, with S extract inducing greater attraction (*χ2* = 12.577, *df* = 1, *P* < 0.05). A similar attractiveness pattern is seen when the S extract is paired with water (*χ2* = 22.5, *df* = 1, *P* < 0.05). When given opportunities to select between water and the sweetened extract, MP workers approach both Arabica infusions, but the latter extract is significantly less attractive than the first one (*χ2* = 16.39, *df* = 1, *P* < 0.05) (Fig. 3C).

## DISCUSSION

The most important observation of this study is that coffee influenced the behavior of *Tapinoma indicum*, *Monomorium pharaonis* and *Solenopsis geminata*. That influence also showed different patterns according to species, coffee concentration, and sugar content.

The fact that coffee elicited attraction to the workers of formicine ants, irrespective of species seems in agreement with an earlier work from Wright et al. (2013), who substantiated the ability of the main coffee ingredient—caffeine to manipulate the olfactory and behavioral responses of hymenopterans. Bees and ants are hymenopterans from the aculeate subclade, a group characterized by an increased level of sociality (Wilson, 1971; Bradley et al., 2009). Besides ecological and behavioral traits, ants and bees are genetically related (Johnson et al., 2013) and have marked similarities in their sensory system and functioning (Gronenberg, 2008). Their antennae are equipped with a variety of types of sensory sensilla, many of which specialized for chemoreceptive functions (Kropf, Kelber, Bieringer, and Rössler, 2014; Barsagade, Tembhare, and Kadu, 2013). In both formicine and honeybees, the olfactory sensilla hold olfactory receptor neurons that are connected to glomeruli, the functional unit of the antennal lobe (Anton and Homberg, 2013; Gronenberg, 2008). This structure is the first relay station for the processing of olfactory information (Zube, et al., 2008). Ants possess over 400 glomeruli (Gronenberg, 2008), representing about twice the number of glomeruli in honeybee (Flanagan and Mercer, 1989). Given this, it is tempting to advocate that all the tested ant species have behaviorally responded to the coffee odor because they all possess coffee odorants-specific olfactory receptors. It is possible that the discrepancies in behavioral response intensities observed between the *T. indicum*, *Monomorium pharaonis,* and *Solenopsis geminata* owe to the differences in glomeruli abundance and/or coffee aroma-mediated olfactory detection capacity. In favor of this insinuation, it has been documented that the number of glomeruli differs across formicine ant species (Goll, 1967; Nishikawa et al., 2008; Zube, et al., 2008). The evidence also exist that demonstrate a close link between glomerulus abundance and smell detection competence (Gronenberg, 2008; Kelber, et al., 2009) as well as the quantity of odors that can be discriminated (Gronenberg, 2008).

Within the tube olfactometer, we showed a major effect of coffee concentration on the behavioral responses of the workers of *T. indicum*, *Monomorium pharaonis* and *Solenopsis geminata*. Differences in behavioral reaction to chemical cues have often been associated with the concentrations of chemical compounds. Youngsteadt et al. (2007) examined the attractiveness of a blend of five compounds from *Peperomia macrostachya* seeds, one component—geranyl linalool from the same seeds and a solvent (control) on the formicine ant, *Camponotus femoratus*. They found that workers preferred the blend and its attractiveness gradually decreased in the order of geranyl linalool > solvent. The abating attraction level seen when proceeding from blend to geranyl linalool to solvent, distinctly indicates an odor concentration-dependent attractiveness. Coffee aroma is perceived via smelling and the perception of coffee smell is dependent upon both the concentration of the compound and its odor threshold (Coffee Research Org, 2016). There are more than 800 volatile aromatic compounds in coffee (Coffee Research Org, 2016) that occur naturally (Illy and Illy, 2015) and during roasting (Farah, 2012). The flavor and aroma of coffee have been well-documented (Clarke, 1985; Blank, et al., 1991). For instance, damascenone and 3-mercapto- 3-methylbutylformate give a honey-like, fruity and catty odor respectively. 3-Methyl-2-buten-1-thiol and 2-Isobutyl-3-methoxypyrazine are responsible for the amine-like and earthy smells. The phenolic, buttery and spicy scents originate from Guaiacol, 2,3-Butanedione and 4-vinylguaiaco accordingly. The methional, vanillin, and furaneol are responsible for the potato-like, vanilla and caramel-like odors. Sotolon and abhexon have a seasoning-like smell.

The present choice experiment was conducted with three different Arabica concentrations: 0.01%, 0.05%, and 0.10%. For all species, workers tended to prefer 0.01% and 0.05% over 0.10%. All these extracts came from Arabica and therefore contain the same chemicals. Hence, the observed differences in the attractiveness level between the low (0.01% and 0.05%) and the highest (0.10%) concentrations may have occurred due to at least two reasons. First, the concentration 0.10% may contain a greater amount of odorous compounds than the two other test concentrations (0.01% and 0.05%), and it is likely that the increased presence of such compounds has produced a strong smell that was repulsive to the workers. A second possible explanation for the discrepancy in the Arabica attraction between the highest and low concentrations could be related to the difference in color stimulus. Optically, the 0.01% concentration had a light brown color, while the 0.10% was dark brown. Formicine ants are known to be capable of differentiating between colors (Camlitepe and Aksoy, 2010) and do not like strong colors (Science Buddies, 2011). It is possible that the dark brown color of the 0.10% Arabica matched the appearance of potential predators i.e., *Anoplolepis gracilipes*—a predator (Konopik, et al., 2014) common in the areas where the experimental workers originated from— and acted as a repulsive signal to workers.

There was an obvious association between the presence of sugar in Arabica extract and the behavioral responses of the tested ants. Significant attractions to sugared Arabica were observed for workers of TI, MP and SG when unsweetened Arabica or water was the other option offered to them. Similar observations were reported previously in formicine ants. Dieng et al. (2016), in an experiment, worked with the black crazy ant exposed its workers to four extracts of tea with different sugar contents and found a significantly greater preference for two sweetest meals over those without sugar or with the least amount of sugar. In related work, Stanley and Robinson (2007), when dealing with *Paratrechina bourbonica* exposed its workers to five baits and reported sugared solutions among the most attractive baits. Meanwhile, Nyamukondiwa and Addison (2014) examined food preferences of the Argentine ant, the black cocktail ant and *Anoplolepis custodiens* in the wild. They observed that their foragers had a preference for sugared water and honey. To shed light on the macronutrient preference and consumption of the formicine *Camponotus pennsylvanicus*, Cannon (1998) used simple sugars with various levels of sweetness. He reported a preference for sucrose and fructose—known to be sweeter than glucose (WordPress.com, 2009). In the sugar effect study, three extracts were used: Arabica with 0.50% sugar or “S”, Arabica without sugar or “NS” and water (Control or C). The extract “S” was generated by dissolving 0.25 g of sugar (also known as table sugar or sucrose (WordPress.com, 2009) in 50 mL of a 0.01% Arabica solution. Generally, formicine ants including TI (*Tapinoma indicum)* (Lee, 2008), MP (*Monomorium pharaonis*) (Sorensen and Vinson, 1981) are naturally highly active (Harris, de Ibarra, Graham, and Collett, 2005). For most social insects including formicine ants, sugar is a ready-to-use energy source (Chong, et al., 2002). In most ant species, sugars have positive effects, supplying energy for various activities (Kay, et al., 2004). Based on these reports, the observed strong attraction of sugar to TI, MP and SG foragers may have been due to the need of energy supply for their foragers. In support of this, it has been reported that active ant workers are constantly in an energy deficient state (Sorensen and Vinson, 1981).

Besides giving insight into the behavioral ecology of three different formicine species, the results of the current study illustrated the potential of coffee, especially Arabica, as a valuable source in baiting strategy against ant pests.

## FUNDING

The authors would like to acknowledge Ministry of Higher Education (MOHE) for funding the research under Fundamental Research Grant (FRGS) (FRGS: 203 / PBIOLOGI / 6711360).

## References

Anon (1986). Pharaoh’s ants. Monomorium pharaonis (L.). National Pest Control Association, ESPC 032155, Dunn Loring, iiiiiiVA, 6 pp.

Anton, S., and Homberg, U. (2013). Antennal lobe structure. In B. S. Hansson (Ed.), Insect olfaction (pp. 98–125). Heidelberg: Springer Science and Business Media.

Barsagade, D. D., Tembhare, D. B., and Kadu, S. G. (2013). Microscopic structure of antennal sensilla in the carpenter ant *Camponotus compressus* (Fabricius) (Formicidae: Hymenoptera). Asian Myrmecology 5, 113–120.

Blank, I., Sen, A., and Grosch, W. (1991). Aroma impact compounds of Arabica and Robusta coffee. Qualitative and quantitative investigations. In Proceedings of the 14th International Conference on Coffee Science (pp. 117–129). San Francisco. Retrieved from <u>http://www.asic-cafe.org/en/proceedings</u> (accessed 13.1.17).

Bradley, T. J., Briscoe, A. D., Brady, S. G., Contreras, H. L., Danforth, B. N., Dudley, R., … Reppert, S. M. (2009). Episodes in insect evolution. Integrative and Comparative Biology 49(5), 590–606.

Camlitepe, Y., and Aksoy, V. (2010). First evidence of fine colour discrimination ability in ants (Hymenoptera, Formicidae). Journal of Experimental Biology 213(1), 72–77.

Cannon, C. A. (1998). Nutritional ecology of the carpenter ant *Camponotus pennsylvanicus* (De Geer): macronutrient preference and particle consumption. Ph.D dissertation, Virginia Polytechnic Institute and State University, Blacksburg, VA.

Chang, V., and Ota, A. K. (1990). Ant control in Hawaiian drip irrigation systems. In R. K. Vander Meer, K. Jaffe and A. Cedeno (Eds.), Applied myrmecology: a world perspective (pp. 708–715). Boulder, CO: Westview Press.

Choe, D. H., Ramírez, S. R., and Tsutsui, N. D. (2012). A silica gel based method for extracting insect surface hydrocarbons. Journal of Chemical Ecology 38(2), 176–187.

Cholakov, G., Toteva, V., Nikolov, R., Uzunova, S., and Yanev, S. (2013). Extracts from coffee by-products as potential raw materials for fuel additives and carbon adsorbents. Journal of Chemical Technology and Metallurgy 48(5), 497–504.

Chong, A., Chong, N. L., Yap, H. H., and Lee, C. Y. (2002). Effects of starvation on nutrient distribution in the Pharaoh ant, *Monomorium pharaonis* (Hymenoptera: Formicidae) workers and various larval stages. In Proceedings of the 4th International Conference on Urban Pests. Virginia, VA: Pocahontas Press, Inc. (pp. 121–127).

Clarke, R. J. (1985). The technology of converting green coffee into the beverage. In M. N. Clifford and K. C. Willson (Eds.), Coffee: Botany, biochemistry and production of beans and beverage (pp. 375–393). Connecticut, CT: The AVI Publishing Company, Inc.

Clarke, R. J., and Macrae, R. (2012). Coffee: Volume 1: Chemistry. New York, NY: Springer Science and Business Media.

Coffee Research Org. 2016. Coffee chemistry: Coffee aroma. Retrieved from http://www.coffeeresearch.org/science/aromamain.htm (accessed 14.1.17)

Coggins, J. M. (2013). Crazy ants that feast on electronics and are invading the U.S cannot be killed with normal insecticide. Retrieved from http://www.dailymail.co.uk/news/article-2338502/Crazy-ants-terrorizing-parts-U-S-resistant-chemicals-kill-species.html(accessed 14.1.17)

Derraik, J. G. B., and Slaney, D. (2005). The toxicity of used coffee grounds to the larvae of *Ochlerotatus (Finlaya) notoscriptus* (Skuse) (Diptera: Culicidae). The Annals of Medical Entomology 14, 14–24.

Dieng, H., Mohd Zawawi, R., Mohamed Yusof, N. I. S., Ahmad, A. H., Abang, F., Abd Ghani, I., … and Noweg, G. T. (2016). Green tea and its waste attract workers of formicine ants and kill their workers—implications for pest management. Industrial Crops and Products 89, 157–166.

Dodgson, L. (2015). Supersniffing ants smell things humans can't. Retrieved from http://www.livescience.com/51843-ants-can-outsniff-humans.html (accessed 14.1.17)

Dumpert, K. (1981). The social biology of ants. New York, NY: Chapman and Hall.

Dupuy, F., Sandoz, J. C., Giurfa, M., and Josens, R. (2006). Individual olfactory learning in Camponotus ants. Animal Behaviour 72(5), 1081–1091.

Farah, A. (2012). Coffee constituents. In Y. F. Chu (Ed.), Coffee: Emerging health effects and disease prevention (pp. 21–58). Iowa, IA: John Wiley and Sons Inc.

Fisk, I. D., Kettle, A., Hofmeister, S., Virdie, A., and Kenny, J. S. (2012). Discrimination of roast and ground coffee aroma. Flavour 1, 14.

Flanagan, D., and Mercer, A. R. (1989). Morphology and response characteristics of neurones in the deutocerebrum of the brain in the honeybee *Apis mellifera*. Journal of Comparative Physiology A 164(4), 483–494.

Goll, W. (1967). Strukturuntersuchungen am Gehirn von Formica. Zeitschrift für Morphologie und Ökologie der Tiere 59(2), 143–210.

Gronenberg, W. (2008). Structure and function of ant (Hymenoptera: Formicidae) brains: Strength in numbers. Myrmecological News, 11, 25–36.

Grosch, W., 1998. Flavour of coffee: A review. Nahrung 42(06), 344–350.

Haack, K. D., and Granovsky, T. A. (1990). Ants. In K. Story and Moreland D (Eds.), Handbook of Pest Control (pp. 415–479). Ohio, OH: Franzak and Foster Co.

Hammond, P. (2011). Atlas of the world’s strangest animals. New York, NY: Marshall Cavendish Reference.

Harris, R. A., de Ibarra, N. H., Graham, P., and Collett, T. S. (2005). Ant navigation: Priming of visual route memories. Nature 438(7066), 302–302.

Harris-Lovett, S. 2015. For ants, order is maintained by odor. Retrieved from http://www.poconorecord.com/article/20150816/NEWS/150819609 (accessed 14.1.17)

Higgins, W., Bell, D., Silcox, C., and Hollbrook, G. (1997). Perimeter baits, spray, or combinations: Which provide longer odorous house ant relief for residential accounts. In Proceedings of the 4th International Conference on Urban Pests (pp. 134–140). Virginia, VA: Pocahantas Press.

Hill, D. S. (1987). Agricultural insect pests of temperate regions and their control. New York, NY: Cambridge University Press.

Hölldobler, T., and Wilson, E. O. (1990). The ants. Massachusetts, MA: Havard University Press.

Illy, E., and Illy, A. (2015). The complexity of coffee. Scientific American 24, 10–15.

John and Sarah. (2010). Interesting facts about ants. Retrieved from http://www.lingolex.com/antsteach.htm (accessed 14.1.17)

Johnson, B. R., Borowiec, M. L., Chiu, J. C., Lee, E. K., Atallah, J., and Ward, P. S. (2013). Phylogenomics resolves evolutionary relationships among ants, bees, and wasps. Current Biology 23(20), 2058–2062.

Kay, A. D., Scott, S. E., Schade, J. D., and Hobbie, S. E. (2004). Stoichiometric relations in an ant‐treehopper mutualism. Ecology Letters 7(11), 1024–1028.

Kelber, C., Rössler, W., Roces, F., and Kleineidam, C. J. (2009). The antennal lobes of fungus-growing ants (Attini): neuroanatomical traits and evolutionary trends. Brain, Behavior and Evolution 73(4), 273–284.

Konopik, O., Gray, C. L., Grafe, T. U., Steffan-Dewenter, I., and Fayle, T. M. (2014). From rainforest to oil palm plantations: Shifts in predator population and prey communities, but resistant interactions. Global Ecology and Conservation 2, 385–394.

Kropf, J., Kelber, C., Bieringer, K., and Rössler, W. (2014). Olfactory subsystems in the honeybee: Sensory supply and sex specificity. Cell and Tissue Research 357(3), 583–595.

Krushelnycky, P. D., and Gillespie, R. G. (2008). Compositional and functional stability of arthropod communities in the face of ant invasions. Ecological Applications 18(6), 1547–1562.

Man, L. S., and Lee, C. Y. (2014). Structure-invading pest ants in healthcare facilities in Singapore. Sociobiology 59(1), 241–249.

Lee, C. Y. (2002). Tropical household ants – pest status, species diversity, foraging behavior and baiting studies. In Proceedings of the 4th International Conference on Urban Pests (pp. 3–18). Virginia, VA: Pocahontas Press.

Lee, C. Y. (2008). Sucrose bait base preference of selected urban pest ants (Hymenoptera: Formicidae). In Proceedings of the 6th International Conference on Urban Pests (pp. 59–63). Veszprém: OOK-Press Kft.

Lucas, J. R., and Invest, J. F. (1993). Factors involved in the successful use of hydramethylnon baits in household and industrial pest control. In Proceedings of the 1st International Conference on Insect Pests in the Urban Environment (pp. 99–106). Cambridge: BPCC Wheatons Exeter.

Nestel, D., and Dickschen, F. (1990). The foraging kinetics of ground ant communities in different Mexican coffee agroecosystems. Oecologia 84(1), 58–63.

Nishikawa, M., Nishino, H., Misaka, Y., Kubota, M., Tsuji, E., Satoji, Y., Ozaki, M., and Yokohari, F. (2008). Sexual dimorphism in the antennal lobe of the ant Camponotus japonicus. Zoological Science 25(2), 195–204.

Nyamukondiwa, C., and Addison, P. (2014). Food preference and foraging activity of ants: Recommendations for field applications of low-toxicity baits. Journal of Insect Science 14, 48.

Osae, M. Y., Cobblah, M. A., Djankpa, F. T., Lodoh, E., and Botwe, P. K. (2011). Development of a bait system for the Pharaoh’s ant, *Monomorium Pharaonis* L.(Hymenoptera: Formicidae). West African Journal of Applied Ecology 18(1), 29–38.

Passera, L. (1994). Characteristics of tramp species. In D. F. Williams (Ed.), Exotic ants: Biology, impact and control of introduced species, (pp. 23–43). Colorado, CO: Westview Press.

Potter, M. F., and Hillery, A. E. (2002). Exterior-targeted liquid termiticides: An alternative approach to managing subterranean termites (Isoptera: Rhinotermitidae) in buildings. Sociobiology 39(3), 373–405.

Prins, A. J. (1985). Formicoidea. In C. H. Scholtz and E. Holm (Eds.), Insects of southern Africa (pp. 443–451). Butterworths.

Roces, F. (1994). Odour learning and decision-making during food collection in the leaf-cutting ant *Acromyrmex lundi*. Insectes Sociaux 41(3), 235–239.

Rust, M. K., and Choe, D. H. (2012). Pest notes: Ants. Retrieved from http://ipm.ucanr.edu/PMG/PESTNOTES/pn7411.html (accessed 14.1.17)

Rust, M. K., Reierson, D. A., and Klotz, J. H. (2002). Factors affecting the performance of bait toxicants for Argentine ants (Hymenoptera: Formicidae). In Proceedings of the 4th International Conference on Urban Pests (pp. 115–120).

Satho, T., Dieng, H., Itam Ahmad, M. H., Ellias, S., Ahmad, A. H., Abang, F., … Nolasco-Hipolito, C. (2015). Coffee and its waste repel gravid Aedes albopictus females and inhibit the development of their embryos. Parasites and Vectors 8(1), 1.

Seo, H. S., Hirano, M., Shibato, J., Rakwal, R., Hwang, I. K., and Masuo, Y. (2008). Effects of coffee bean aroma on the rat brain stressed by sleep deprivation: a selected transcript-and 2D gel-based proteome analysis. Journal of Agricultural and Food Chemistry 56(12), 4665–4673.

Science Buddies. (2011). Color-soaked bugs: Color sense in insects. Retrieved from http://www.sciencebuddies.org/blog/2011/09/color-soaked-bugs-color-sense-in-insects.php (accessed 14.1.17)

Smith, E. H., and Whitman, R. C. (1992). Field guide to structural pests. Virginia, VA: National Pest Control Association.

Sorensen, A. A., and Vinson, S. B. (1981). Quantitative food distribution studies within laboratory colonies of the imported fire ant, *Solenopsis invicta* Buren. Insectes Sociaux 28(2), 129–160.

Stanley, M. C. (2004). Review of the efficacy of baits used for ant control and eradication. Landcare research contract report: LC0404/044. Retrieved from http://www.landcareresearch.co.nz/research/biocons/invertebrates/ants/BaitEfÞcacyReport.pdf (accessed 14.1.17)

Stanley, M. C., and Robinson, W. A. (2007). Relative attractiveness of baits to Paratrechina longicornis (Hymenoptera: Formicidae). Journal of Economic Entomology 100(2), 509–516.

Shirwaikar, A., Issac, D., and Malini, S. (2004). Effect of Aerva lanata on cisplatin and gentamicin models of acute renal failure. Journal of Ethnopharmacology 90(1), 81–86.

WordPress.com. (2009). Simple sugars: Fructose, glucose and sucrose. Retrieved from https://cdavies.wordpress.com/2009/01/27/simple-sugars-fructose-glucose-and-sucrose (accessed 14.1.17)

Williams, D. F., Lofgren, C. S., and Vander Meer, R. K. (1990). Fly pupae as attractant carriers for toxic baits for red imported fire ants (Hymenoptera: Formicidae). Journal of Economic Entomology 83(1), 67–73.

Wilson, W. T. (1971). Resistance to American foulbrood in honey bees. XI: fate of Bacillus larvae spores ingested by adults. Journal of Invertebrate Pathology 17(2), 247–255.

Wilson, E. O. (2005). Environment: Early ant plagues in the New World. Nature 433(7021), 32–32.

Wolf, H., and Wehner, R. (2000). Pinpointing food sources: olfactory and anemotactic orientation in desert ants, *Cataglyphis fortis*. Journal of Experimental Biology 203(5), 857–868.

Wright, G. A., Baker, D. D., Palmer, M. J., Stabler, D., Mustard, J. A., Power, E. F., Borland, A. M., and Stevenson, P. C. (2013). Caffeine in floral nectar enhances a pollinator's memory of reward. Science 339(6124), 1202–1204.

Youngsteadt, E. (2007). Complex chemical recognition mediates an ant-seed mutualism in the Amazonian rainforest. In The 2007 ESA Annual Meeting, December 9–12, 2007.

Yusuf, A. A., Crewe, R. M., and Pirk, C. W. (2014). Olfactory detection of prey by the termite-raiding ant *Pachycondyla analis*. Journal of Insect Science 14(1), 53.

Zube, C., Kleineidam, C. J., Kirschner, S., Neef, J., and Rössler, W. (2008). Organization of the olfactory pathway and odor processing in the antennal lobe of the ant *Camponotus floridanus*. Journal of Comparative Neurology 506(3), 425–441.

